# The impact of DNA polymerase and number of rounds of amplification in PCR on 16S rRNA gene sequence data

**DOI:** 10.1101/565598

**Authors:** Marc A Sze, Patrick D Schloss

**Affiliations:** Department of Microbiology and Immunology, University of Michigan, Ann Arbor, MI

## Abstract

PCR amplification of 16S rRNA genes is a critical, yet under appreciated step in the generation of sequence data to describe the taxonomic composition of microbial communities. Numerous factors in the design of PCR can impact the sequencing error rate, the abundance of chimeric sequences, and the degree to which the fragments in the product represent their abundance in the original sample (i.e. bias). We compared the performance of high fidelity polymerases and varying number of rounds of amplification when amplifying a mock community and human stool samples. Although it was impossible to derive specific recommendations, we did observe general trends. Namely, using a polymerase with the highest possible fidelity and minimizing the number of rounds of PCR reduced the sequencing error rate, fraction of chimeric sequences, and bias. Evidence of bias at the sequence level was subtle and could not be ascribed to the fragments’ fraction of bases that were guanines or cytosines. When analyzing mock community data, the amount that the community deviated from the expected composition increased with rounds of PCR. This bias was inconsistent for human stool samples. Overall the results underscore the difficulty of comparing sequence data that are generated by different PCR protocols. However, the results indicate that the variation in human stool samples is generally larger than that introduced by the choice of polymerase or number of rounds of PCR.

**Importance:** A steep decline in sequencing costs drove an explosion in studies characterizing microbial communities from diverse environments. Although a significant amount of effort has gone into understanding the error profiles of DNA sequencers, little has been done to understand the downstream effects of the PCR amplification protocol. We quantified the effects of the choice of polymerase and number of PCR cycles on the quality of downstream data. We found that these choices can have a profound impact on the way that a microbial community is represented in the sequence data. The effects are relatively small compared to the variation in human stool samples, however, care should be taken to use polymerases with the highest possible fidelity and to minimize the number of rounds of PCR. These results also underscore that it is not possible to directly compare sequence data generated under different PCR conditions.

## Introduction

16S rRNA gene sequencing is a powerful and widely used tool for surveying the structure of microbial communities (1–3). This approach has exploded in popularity with advances in sequencing throughput such that it is now possible to characterize numerous samples with thousands of sequences per sample. Many factors can impact how a natural community is represented by the sequencing data including the method of acquiring samples (4–8), storage conditions (4–6, 9–12), extraction methods (13), amplification conditions (8, 14, 15), sequencing method (15–17), and analytical pipeline (15, 18–20). The increased sampling depth that is now available relative to previous Sanger sequencing-based methods is expected to compound the impacts of an investigator’s choices and the interpretation of their results.

One step in the generation of 16S rRNA gene sequence data that has been long known to have a significant impact on the description of microbial communities is the choice of conditions for PCR amplification (8, 14, 15). Factors such as the choice of primers have an obvious impact on which populations will be amplified (18, 21). However, a variety of PCR artifacts can also impact the perception of a community including the formation of chimeras (14, 22–24), misincorporation of nucleotides (23, 25, 26), preferential amplification of some populations over others leading to bias (24, 27–33), and accumulation of random amplification events leading to PCR drift (24, 27, 32, 34). Many bioinformatic tools have been developed to identify chimeras; however, there are significant sensitivity and specificity tradeoffs (14, 35). Laboratory-based solutions to minimize chimera formation have also been proposed such as minimizing the amount of template DNA in the PCR, minimizing the number of rounds of PCR, minimizing the amount of shearing in the template DNA, using DNA polymerases that have a proof-reading ability, and emulsion PCR (14, 23, 36). Others have attempted to account for PCR bias using modeling approaches (29, 37). In cases where such modeling approaches have been successful, it has been with relatively small communities with consistent composition (29). To minimize PCR drift, some investigators pool technical replicate PCRs hoping to average out the drift (34). Other factors that have been shown to impact the formation of PCR artifacts are outside the control of an investigator including the fraction of DNA bases that are guanines or cytosines, the variation in the length of the targeted region across the community, the sequence of the DNA that flanks the template, and the genetic diversity of the community (28, 30–33). Early investigations of the factors that lead to the formation of PCR artifacts focused on analyzing binary mixtures of genomic DNA and 16S rRNA gene fragments to explore PCR biases and chimera formation. Although these studies were instrumental in forcing researchers to be cautious about the interpretation of their results, we have a poor understanding of how these factors affect the formation of PCR artifacts in more complex communities.

The influence that the choice of DNA polymerase has on the formation of PCR artifacts has not been well studied. There has been recent interest in how the choice of the hypervariable region and data analysis pipelines impact the sequencing error rate (15, 18–20); however, these studies use the same DNA polymerase in the PCR step and implicitly assume that the rate of nucleotide misincorporation from PCR are significantly smaller than those from the sequencing phase. There has been more limited interest in the impact that DNA polymerase choice has on the formation of chimeras (23, 38). A recent study found differences in the number of OTUs and chimeras between normal and high fidelity DNA polymerases (38). The authors of the study reduced the difference between two polymerases by optimizing the annealing and extension steps within the PCR protocol (38). Yet this optimization was specific for the community they were analyzing (i.e. captive and semi-captive red-shanked doucs) and assumed that if the two polymerases generate the same community structure that the community structure was correct. In fact, the community structure generated by both methods was not free of artifacts, but likely had the same artifacts. A challenge in these types of experiments is having *a priori* knowledge of the true community representation. A mock community with known composition allows researchers to quantify the sequencing error rate, fraction of chimeras, and bias (19); however, mock communities have a limited phylogenetic diversity relative to natural communities. Natural communities, in contrast, have an unknown community composition making absolute measurements impossible. They can be used to validate results from mock communities and to understand the relative impacts of artifacts on the ability to differentiate biological and methodological sources of variation. Given the large number of DNA polymerases available to researchers, it is unlikely that a specific recommendation is possible. Rather, the development of general best practices and understanding the impact of PCR artifacts on an analysis are needed.

This study investigated the impact of choice of high fidelity DNA polymerase and the number of rounds of amplification on the formation of PCR artifacts using a mock community and human stool samples. It was hypothesized that additional rounds of PCR would exacerbate the number of artifacts. We tested (i) the effect of the polymerase on the error rate of the bases represented in the final sequences, (ii) the fraction of sequences that appeared to be chimeras and the ability to detect those chimeras, (iii) the bias of preferentially amplifying one fragment over another in a mixed pool of templates, and (iv) inter-sample variation in community structure of samples amplified with the same polymerase across the amplification process. To characterize these factors we sequenced a mock community of 8 organisms with known sequences and community structure and human fecal samples with unknown sequences and community structures. We sequenced the V4 region of the 16S rRNA genes from a mock community by generating paired 250 nt reads on the Illumina MiSeq platform. This region and sequencing approach was used because it has been shown to result in a relatively low sequencing error rate and is a widely used protocol (18). To better understand the impact of DNA polymerase choice on PCR artifacts, we selected five high fidelity DNA polymerases and amplified the communities using 20, 25, 30, and 35 rounds of amplification. Collectively, our results suggest that the number of rounds and to a lesser extent the choice of DNA polymerase used in PCR impact the sequence data. The effects are consistent and are smaller than the biological differences between individuals.

## Results

### Sequencing errors vary by the number of cycles and the DNA polymerase used in PCR

The presence of sequence errors can confound the ability to accurately classify 16S rRNA gene sequences and group sequences into operational taxonomic units (OTUs). More importantly, sequencing errors themselves can alter the representation of the community. Therefore, it is important to minimize the number of sequencing errors. Using a widely-used approach that generates the lowest reported error rate, we quantified the error rate by sequencing the V4 region of the 16S rRNA genes from an 8 member mock community. We also removed any contigs that were at least three bases more similar to a chimera of two references than to a single reference sequence (18, 19, 39). Regardless of the polymerase, the error rate increased with the number of rounds of amplification (Figure 1). Using 30 rounds of PCR is a common approach across diverse types of samples. Among the data generated using 30 rounds of PCR the Accuprime polymerase had the highest error rate (i.e. 0.124%) followed by the Platinum (i.e. 0.094%), Phusion (i.e. 0.064%), KAPA (i.e. 0.062%), and Q5 (i.e. 0.060%) polymerases (Figure 1). When we applied a pre-clustering denoising step, which merged the counts of reads within 2 nt of a more abundant sequence (19), the error rates dropped considerably such that the Platinum polymerase had the highest error rate (i.e. 0.014%) followed by the Accuprime (i.e. 0.012%), Q5 (i.e. 0.0053%), Phusion (i.e. 0.0049%), and KAPA (i.e. 0.0049%) polymerases (Figure 1). Although specific recommendations are difficult to make because the phylogenetic diversity of the initial DNA template is likely to have an impact on the results, it is clear that using as few PCR cycles as necessary and a polymerase with the lowest possible error rate is a good guide to minimizing the impact of polymerase on the error rate.

**Figure 1.**
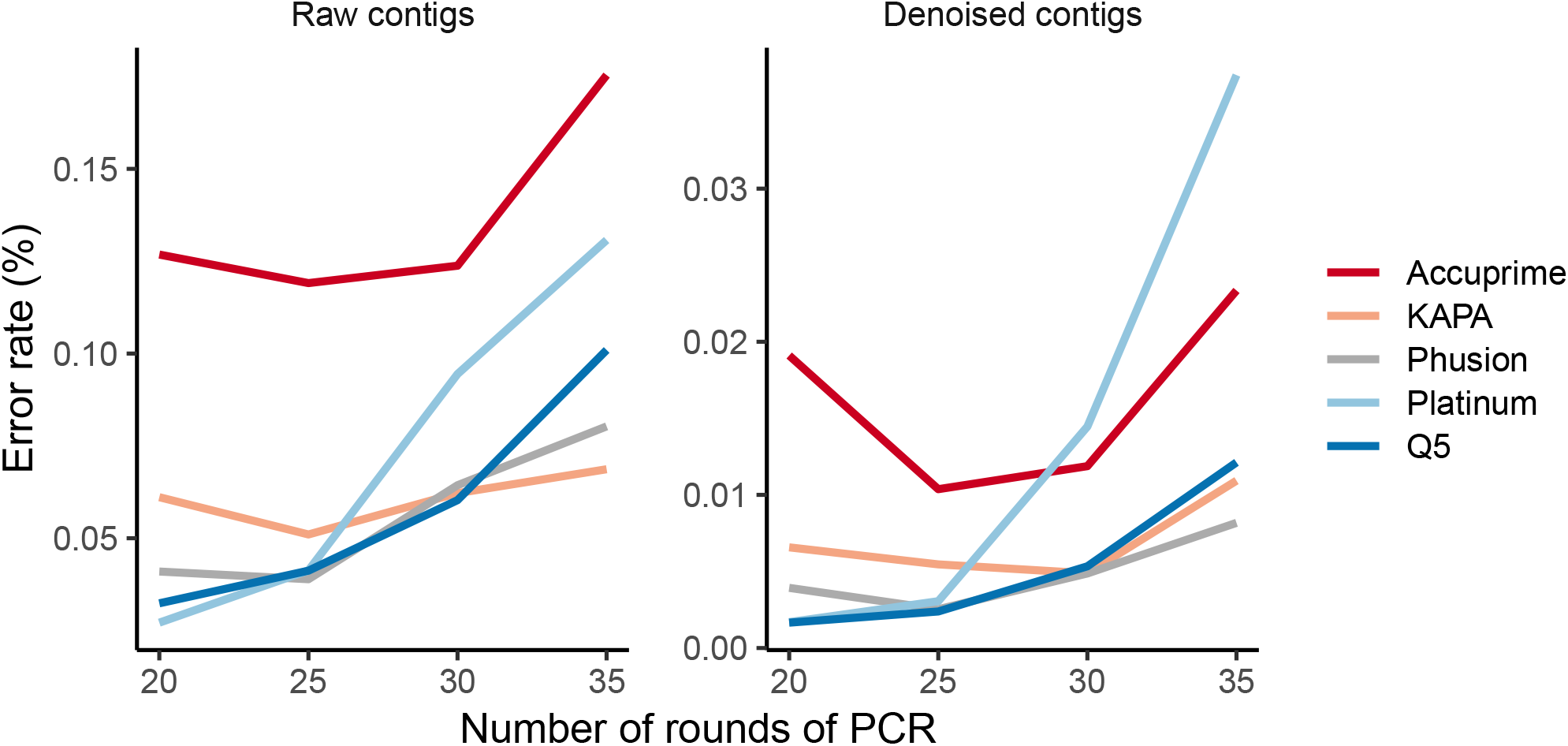
The error rate of assembled mock community sequence reads increases with the number of rounds of PCR; however, much of this error was eliminated by denoising and followed the relative error rates provided by the manufacturers. Each line represents the mean of four replicates.

### The fraction of sequences identified as being chimeric varies by the number of cycles and the DNA polymerase used in PCR

Chimeric PCR products can significantly confound downstream analyses. Although numerous bioinformatic tools exist to identify and remove chimeric sequences with high specificity, their sensitivity is relatively low and can be reduced by the presence of sequencing errors (14, 35). Because the true sequences of the organisms in the mock community were known, we generated all possible chimeras between pairs of V4 16S rRNA gene fragments and used these possible chimeric sequences to screen the sequences generated under the different PCR conditions to detect chimeras. The number of chimeras increased with rounds of amplification (Figure 2A). Interestingly, the fraction of chimeric sequences from the mock community varied by the type of polymerase used. After 30 rounds of PCR, the Platinum polymerase had the highest chimera rate (i.e. 18.2%) followed by the Q5 (i.e. 8.1%), Phusion (i.e. 7.5%), KAPA (i.e. 2.3%), and Accuprime (i.e. 0.9%) polymerases. To explore the characteristics of the chimeras further, we analyzed those chimeras formed after 35 cycles. Because of the uneven number of chimeras generated across the five polymerases, we subsampled the frequency of the chimeras to have the same number of chimeras per polymerase the Q5, Phusion, Accuprime, and Platinum polymerases; the chimeric sequence yield with the KAPA polymerase was significantly lower than the other polymerases and was omitted from our initial comparison. As has been shown previously (14), chimera formation was not random. Among the chimeras that were generated in mock community samples, 4.4% of the chimeras were found across all four polymerases. These chimeras represented between 67.6 and 74.5% of the chimeras generated with each polymerase; they represented 40.4% of the chimeric sequences generated using the KAPA polymerase. These results indicate that the mechanisms leading to the formation of chimeras are largely independent of the properties of the polymerase, but are more likely due to the properties of the sequences.

**Figure 2.**
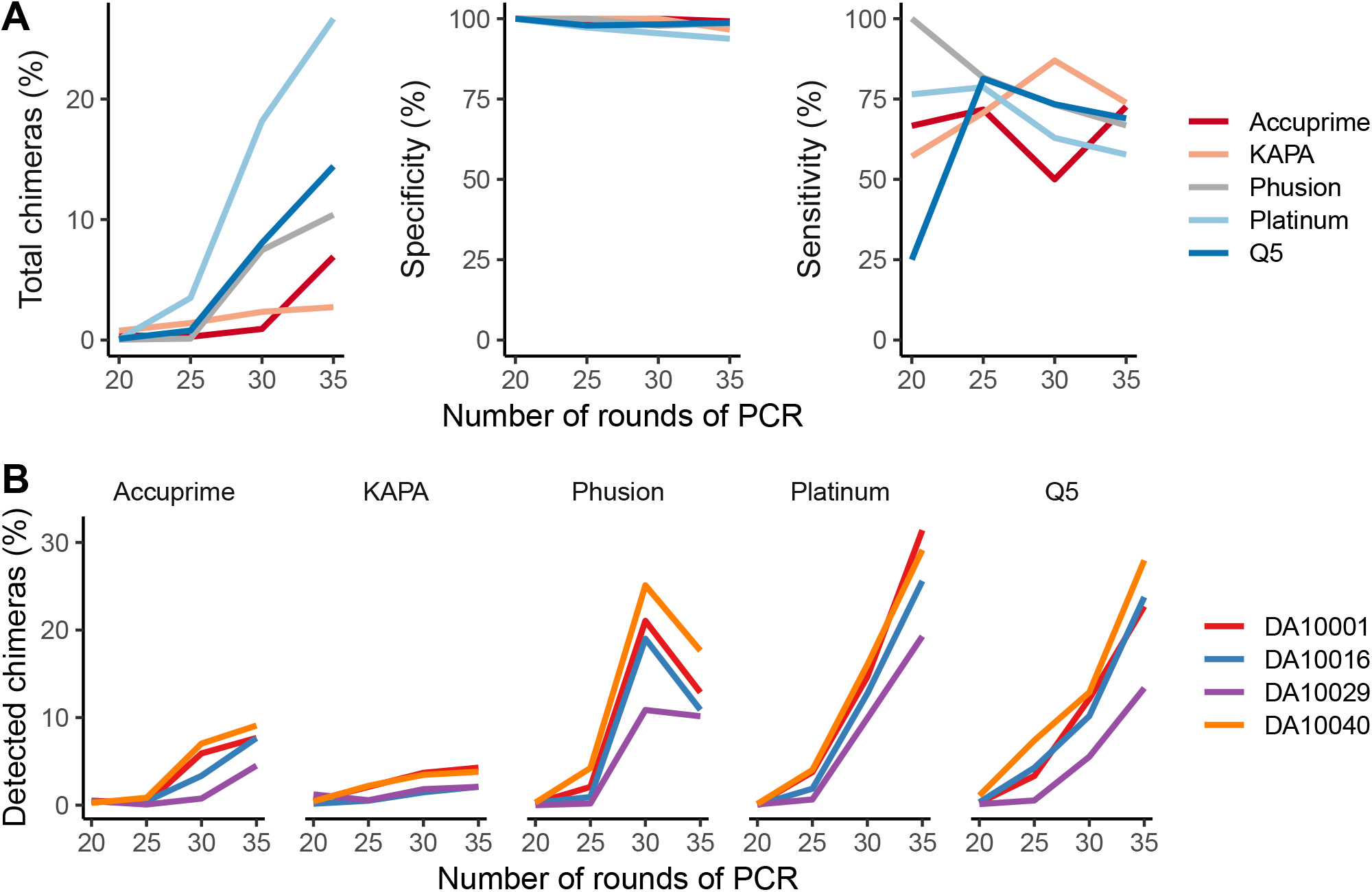
The fraction of all denoised sequences that were identified as being chimeric increases with the number of rounds of PCR used and varied between polymerases. (A) Sequencing of a mock community allowed us to identify the total fraction of sequences that were chimeric as well as the specificity and sensitivity of UCHIME to detect those chimeras. Each line represents the mean of four replicates. (B) Sequencing of four human stool samples after using one of five different polymerases again demonstrated increased rate of chimera formation with increasing number of rounds of PCR and variation across polymerases.

Because our chimera screening procedure could only be applied to mock communities, we used the UCHIME algorithm to model the chimera screening approach that is used in most sequence curation pipelines. By comparing the output of UCHIME to our approach of screening for chimeras using all possible chimeras generated from the mock community sequences, we were able to calculate the UCHIME’s sensitivity and specificity (Figure 2A). The specificity for all polymerases was above 95.4% and showed a weak association with the number of cycles used (Figure 2A). There was considerable inter-polymerase and inter-round of amplification variation in the sensitivity of UCHIME to detect the chimeras from the mock community. This suggested that the residual error rate after pre-clustering the sequence data did not compromise the sensitivity of UCHIME to detect chimeras. The sensitivity of UCHIME varied between 50 and 87.0% when at least 25 cycles were used. The generalizability of these results is limited because we used a single mock community with limited genetic diversity. Although we did not know the true chimera rate for our four human stool samples, we were able to calculate the fraction of sequences that UCHIME identified as being chimeric (Figure 2B). These results followed those from the mock communities: additional rounds of amplification significantly increased the rate of chimeras and there was variation between the polymerases that we used. Although it was not possible to identify the features of a polymerase that resulted in higher rates of chimeras, it is clear that using the smallest number of PCR cycles possible will minimize the impact of chimeras.

### At the sequence level, PCR amplification bias is subtle

Since researchers began using PCR to amplify 16S rRNA gene fragments there has been concern that amplifying fragments from a mixed template pool could lead to a biased representation in the pool of products and would confound downstream analyses (24, 27–33). The mock community was generated by mixing equal amounts of genomic DNA from 8 bacteria resulting in uneven representation of the *rrn* operons across the bacteria as each bacterium had a different genome size and varied in the number of operons in its genome. The vendor of the mock community subjects each lot of genomic DNA to shotgun sequencing to more accurately quantify the actual abundance of each organism in the community. It should be noted that this approach to quantifying abundance is also not without its own biases (40), but does provide an alternative approach to characterizing the structure of the mock community. We compared the vendor reported relative abundance of the 16S rRNA genes from each bacterium in the mock community to the data we generated across rounds of amplification and polymerase (Figure 3). Interestingly, for some bacteria, their representation became less biased with additional rounds of PCR (e.g. *L. fermentum*), while others became more biased (e.g. *E. faecalis*), and others had little change (e.g. *B. subtilis*). Contrary to prior reports (28), the percentage of bases in the V4 region that were guanines or cytosines was not predictive of the amount of bias. Across the strains there was no variation in the length of their V4 regions and they each had the same sequence in the region that the primers annealed. One of the bacteria represented in the mock community, *S. enterica*, had 6 identical copies of the V4 region and 1 copy that differed from those by one nucleotide. The dominant copy had a thymidine and the rare copy had a guanine. We used the sequence data to calculate the ratio of the dominant to rare variants from *S. enterica* expecting a ratio near 6 (Figure S1). The Accuprime, Phusion, Platinum, and Q5 polymerases converged to a ratio of 5.4; however, the ratio for the KAPA polymerase was above 6 for all rounds of PCR (6.1-7.4) and the ratio for Q5 was below 6 for all rounds of PCR (5.3-5.5). Given the subtle nature of the variation in the relative abundances of each 16S rRNA gene fragment, it was not possible to create generalizable rules that would explain the bias.

**Figure 3.**
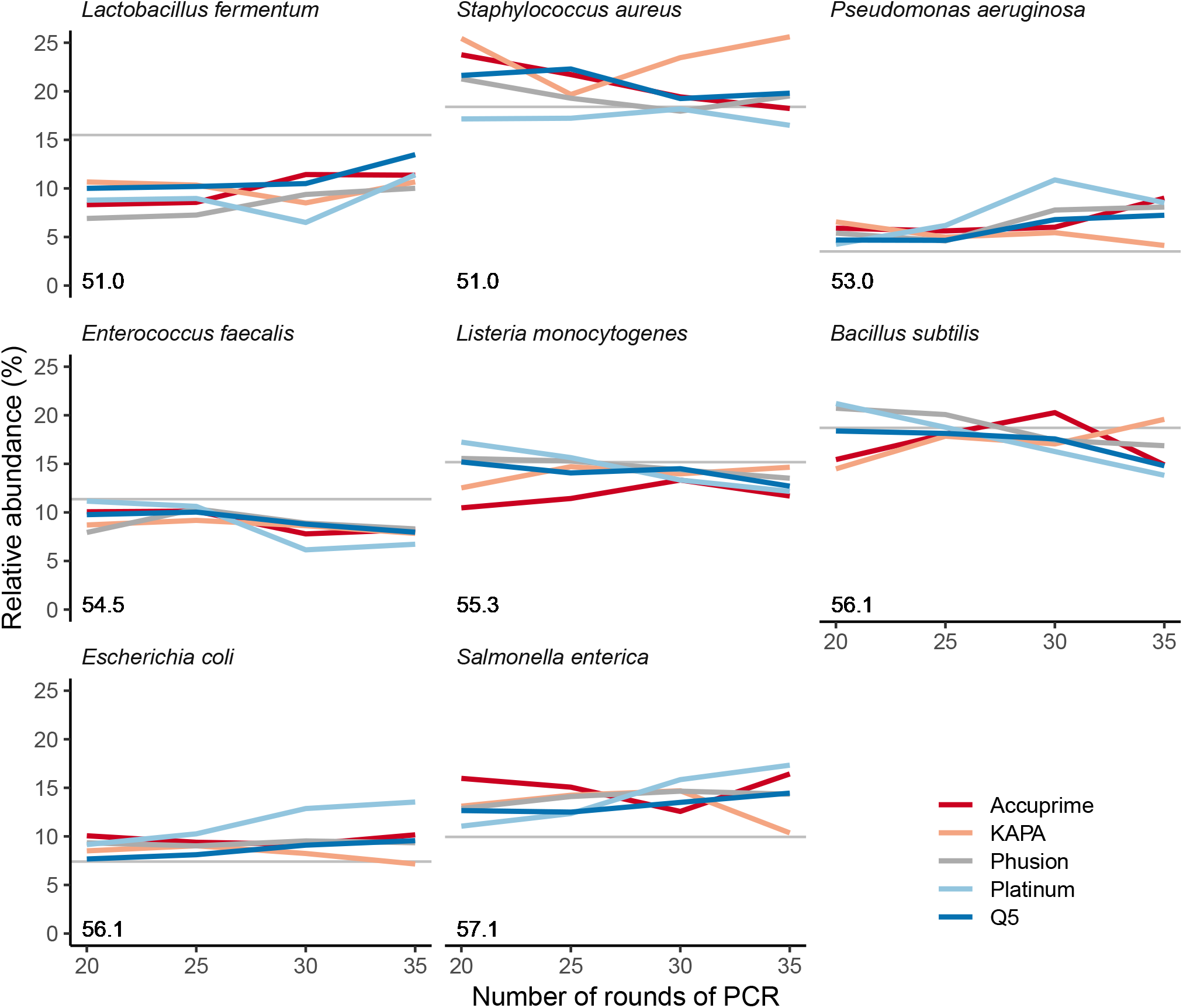
The relative abundances of mock community sequence reads mapped to reference sequences differed subtly from the expected relative abundances as determined by shotgun metagenomic sequencing. Bias did not increase with number of rounds of PCR or vary by polymerase or the guanine and cytosine content of the fragment. The expected relative abundance of each organism is indicated by the horizontal gray line. The percentage of bases that were guanines or cytosines within the V4 region of the 16S rRNA genes in each organism is indicated by the number in the lower left corner of each panel. Each line represents the mean of four replicates.

### At the community level, the effects of PCR amplification bias grow with additional rounds of PCR

Because the variation in bias between polymerases and across rounds of PCR could be artificially inflated due to sequencing errors and chimeras, we analyzed the alpha and beta diversity of the mock community data at different phases of the sequence curation pipeline (Figure 4). First, we removed the chimeras from the mock community data as described above and mapped the individual reads to the OTUs that the 16S rRNA gene fragments would cluster into if there were no sequencing errors. This gave us a community distribution that reflected the distribution following PCR without any artifacts (Figure 4A; “No errors or chimeras”). Although the richness did not change, the Shannon diversity increased with the number of rounds of PCR for all polymerases except the KAPA polymerase, for which the diversity decreased. These data suggest that PCR had the effect of making the community distribution more even than it was originally, except for the data generated using the KAPA polymerase where the evenness decreased. Next, we used the observed sequence errors, but removed chimeras by comparing sequences to all possible chimeras between mock community sequences, and clustered the reads to OTUs (Figure 4A; “Residual errors, complete chimera removal”). The richness and diversity metrics trended higher with higher error rates and number of rounds of PCR. Finally, we used the observed sequence data and the UCHIME algorithm to identify chimeras (Figure 4A; “Residual errors, chimera removal with VSEARCH”). Again, the richness and diversity metrics trended higher with higher error rates and number of rounds of PCR. These comparisons demonstrated that although the bias at the sequence level was subtle, PCR introduces bias at the community level that is exacerbated by errors and chimeras when sequences are clustered into OTUs. When we measured the Bray-Curtis distance between the communities observed after 25 rounds of amplification and those at 30 and 35, distances between 25 and 35 rounds were higher than between 25 and 30 rounds for each of the polymerases by an average of 0.022 units (Figure 4B). The Platinum polymerase varied the most across rounds of amplification (25 vs 30 rounds: 0.13; 25 vs 35 rounds: 0.15). For any number of cycles, the median Bray-Curtis distance between polymerases ranged between 0.074 and 0.11. Although the distances between samples were small, the ordination of these distances showed a clear change in community structure with increasing rounds of PCR (Figure 4C). This observation was supported by our statistical analysis, which revealed that the effect of the number of rounds of PCR (R^2^=0.21, P<0.001) was comparable to the choice of polymerase (R^2^=0.20, P<0.001). These results demonstrate that subtle differences in relative abundances can have an impact on overall community structure. This variation underscores the importance of only comparing sequence data that have been generated using the same PCR conditions.

**Figure 4.**
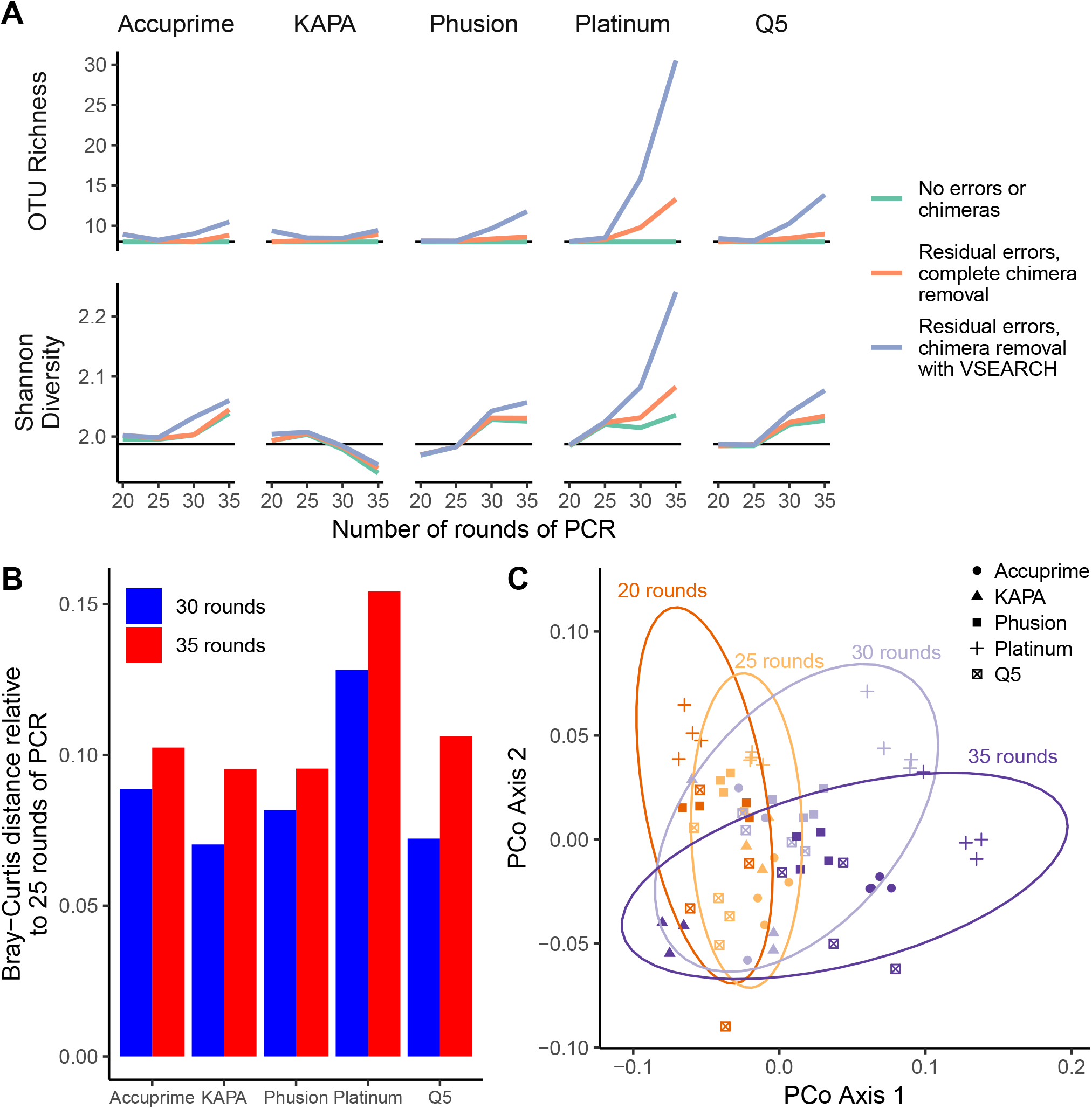
Despite evidence of subtle PCR bias at the genome level, there was significant evidence of bias using community-wide metrics that grew with the number of rounds of PCR when using a mock community. (A) With the exception of the KAPA polymerase data, the richness and Shannon diversity values increased with number of rounds of PCR and the inclusion of residual sequencing errors and chimeras. The horizontal black line indicates the expected richness and diversity. (B) Relative to the mock community sampled after 25 rounds of PCR, the distance to the communities sampled after 30 and 35 rounds of PCR increased for all polymerases. (C) The variation between samples demonstrated a significant change in the community driven by the number of rounds of PCR and the polymerase used. The ellipses represent bivariate normally distributed 95% confidence intervals. The data in A and B represents the mean of four replicates.

### The choice of polymerase or the number of rounds of amplification have little impact on the relative interpretation of community-wide metrics of diversity

We expected that the biases that we observed at the population and community levels using mock community data would be small relative to the expected differences between biological samples. To study this further, we calculated alpha and beta-diversity metrics using the human stool samples for each of the polymerases and rounds of amplification. We calculated the number of observed OTUs and Shannon diversity for each condition and stool sample (Figure 5A). Although there were clear differences between PCR conditions, the relative ordering of the stool samples did not meaningfully vary across conditions. When we characterized the variation between rounds of amplification using human stool samples, the distance between the 25 and 30 rounds and 25 and 35 rounds varied considerably between samples and polymerases (Figure 5B). In general the inter-round variation was lowest for the data generated using the KAPA and Accuprime polymerases. The data generated using the Platinum polymerase was consistent across rounds, but overall, it was more biased than the other polymerases. Considering the average distance across the four samples varied between 0.39 and 0.56, regardless of the polymerases and number of rounds of amplification, any bias due to amplification is unlikely to obscure community-wide differences between samples. In support of this was our principle coordinates analysis of the Bray-Curtis distances, which revealed distinct clusters by stool sample (Figure 5C). Within each cluster there were no obvious patterns related to the polymerase or number of rounds of PCR. Our statistical analysis revealed statistically significant differences in the community structures with the stool sample explaining the most variation (R^2^=0.79, P<0.001), followed by the number of rounds of PCR (R^2^=0.044, P<0.001) and the choice of polymerase (R^2^=0.033, P<0.001). Together, these results indicate that for a coarse analysis of communities, the choice of number of rounds of amplification or polymerase are not important, but that they must be consistent across samples. It is difficult to develop a specific recommendation based on the level of bias across rounds of PCR or polymerases; however, the general suggestion is to use as few rounds of amplification as possible.

**Figure 5.**
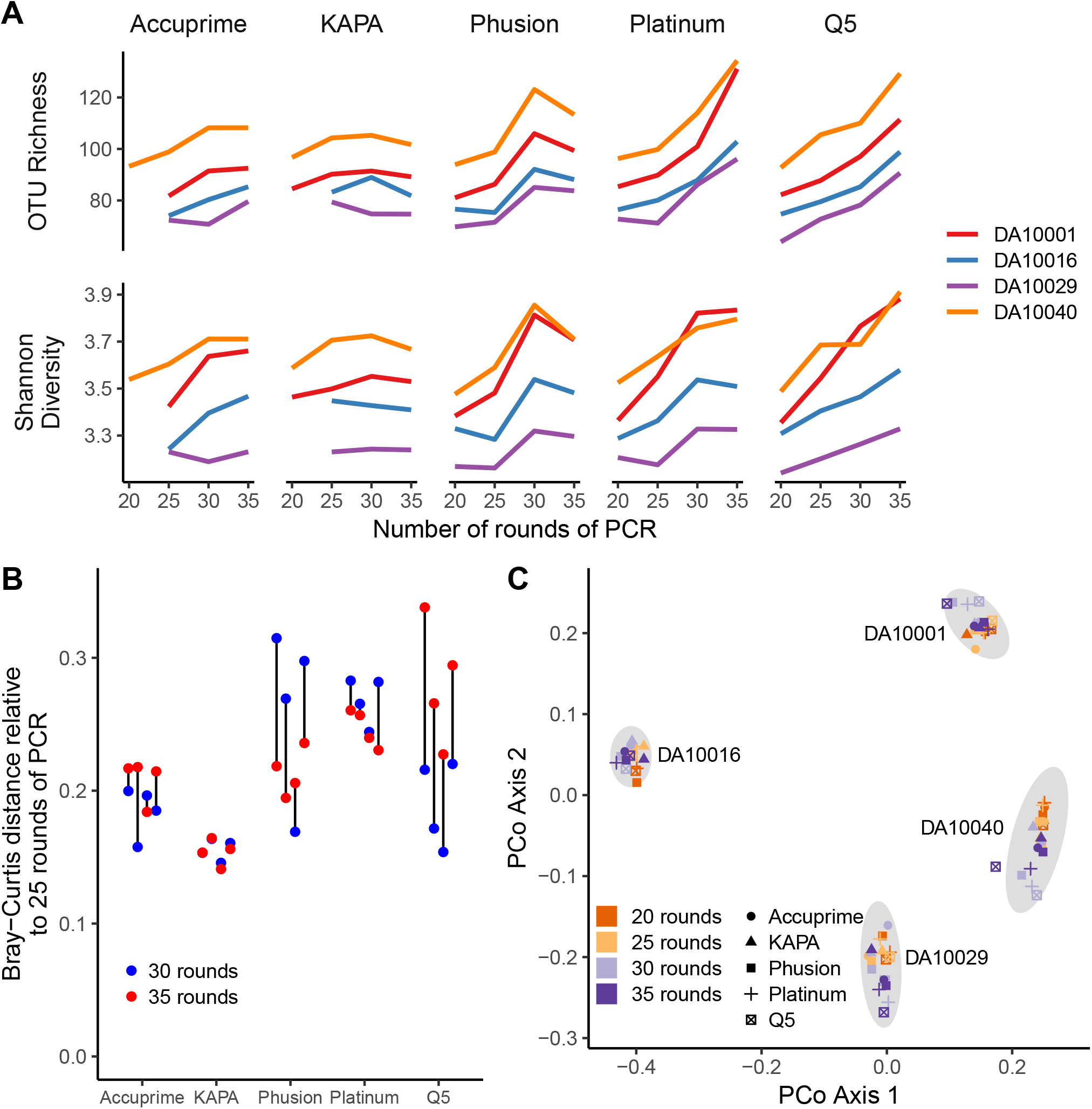
Sequencing of human stool samples indicated clear increase in bias with number of rounds of PCR, however, the bias appeared to be consistent within each sample. (A) With the exception of data collected using the KAPA polymerase, the richness and Shannon diversity values increased with number of rounds of PCR. (B) Relative to the stool communities sampled after 25 rounds of PCR, the distance to the stool communities sampled after 30 and 35 rounds of PCR was inconsistent and there was little difference in variation for data collected using the KAPA polymerase. (C) The variation between stool samples was larger than the amount of variation introduced by varying the number of rounds of PCR or polymerase. The ellipses represent bivariate normally distributed 95% confidence intervals. Results for some samples at 20 cycles are not presented because it was not possible to obtain a sufficient number of reads for those polymerases.

### There is little evidence of a relationship between polymerase or number of rounds of amplification on PCR drift

There have been concerns that the same template DNA subjected to the same PCR conditions could result in different representations of communities because of random drift over the course of PCR. To test this, we determined the average Bray-Curtis distance between replicate reactions using the same polymerase and number of rounds of amplification (Figure 6). Using the mock community data there were no obvious trends. The average Bray-Curtis distance within a set of conditions varied by 0.062 to 0.11 units. Although we did not generate technical replicates of each of the stool samples, the inter-sample variation for each set of conditions was consistent and varied between 0.50 and 0.56 units. These data suggest that amplicon sequencing is robust to random variation in amplification and that any differences are likely to be smaller than what is considered biologically relevant.

**Figure 6.**
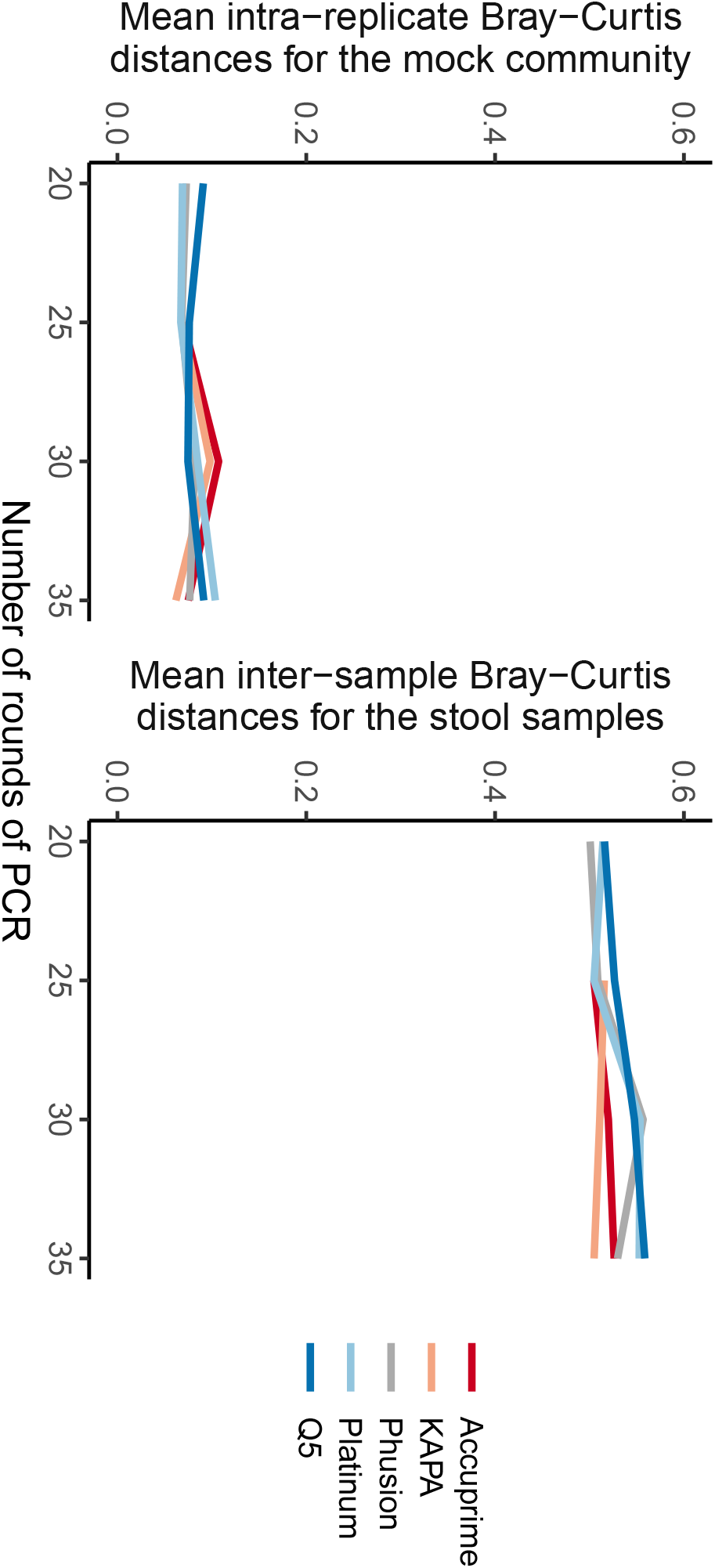
**The average distance between replicates of sequencing the same mock community or between the human stool samples (i.e. drift) did not vary by number of rounds of PCR or by polymerase.**

## Discussion

Our results suggest that the number of rounds of PCR and to a lesser degree the choice of DNA polymerase impact the analysis of 16S rRNA gene sequence data from bacterial communities. Although it was not possible to make direct connections between PCR conditions and specific sources of bias, we were able to identify general recommendations that reduce the amount of error, chimera formation, and bias. Researchers should strive to minimize the number of rounds of PCR and should use a high fidelity polymerase. Although specific PCR conditions impact the precise interpretation of the data, the effects were consistent and were smaller than the biological differences between the samples we tested. Based on these observations, amplicons must be generated by consistent protocols to yield meaningful comparisons. When comparing across studies, values like richness, diversity, and relative abundances must be made in relative and not absolute terms. Furthermore, care must be taken to not directly compare or pool samples from different studies. Instead, it is important to statistically model the study-based variation as has been done in recent meta-analyses that compared relative effect sizes or pooled data using a mixed effects statistical model (41, 42).

The observed sequencing error rates and alpha diversity metrics followed the manufacturers’ measurements of their polymerases’ fidelity (Figure 1). Accuprime and Platinum have fidelity that are approximately 10-times higher than that of Taq whereas the fidelity of Phusion, Q5, and KAPA are more than 100 times higher. Among these polymerases, the KAPA polymerase consistently resulted in a lower error rate, lower chimera rate, and lower bias across rounds of PCR for the mock community samples. Furthermore, among the human samples, the KAPA polymerase consistently had the lowest detected chimera rate and inter-cycle bias. These benefits were most accentuated at 35 cycles. However, in our experience and despite efforts to optimize the yield with the KAPA polymerase, the reactions typically had a high proportion of primer-dimer products and low yield of correctly-sized products. Although the error rate with the Accuprime polymerase was not as low as that with KAPA, we consider it to be an acceptable alternative. Considering polymerase development is an active area of commercial development with potential new polymerases becoming available, it is important for researchers to understand how changing the polymerase impacts downstream analyses for their type of samples.

Over the past 20 years, a large literature has attempted to document various PCR biases and underscored the fact that data based on amplification of DNA from a mixed community are not a true representation of the actual community. In addition to obvious biases imposed by primer selection, other factors inherent in PCR can influence the representation of communities. Factors that can lead to preferential amplification of one fragment over another have included guanine and cytosine composition, length, flanking DNA composition, amount of DNA shearing, and number of rounds of PCR (24, 27–33). These factors may become exacerbated if PCR is performed on multiple samples that vary in their concentration (43). In addition, environmental and reagent contaminants can also have a significant impact on the analysis of low biomass samples (44). Less well understood is the effect of phylogenetic diversity on bias and chimera formation. Communities with low phylogenetic diversity may be more prone to chimera formation since chimeras are more likely to form among closely related sequences (14, 35). The interaction of these various influences on PCR artifacts are complex and difficult to tease apart. Minimizing the level of DNA shearing, controlling for template concentration across samples, and using the fewest number of rounds of PCR with a polymerase that has the highest possible fidelity are strategies that can be employed to minimize the formation of chimeras. Although care should always be taken when choosing a polymerase for 16S rRNA gene sequencing, our observations show that variation among polymerases is smaller than the actual biological variation in fecal communities between individuals.

Even with these strategies it is impossible to remove all PCR artifacts. Beyond the imperfections of the best polymerases, sometimes difficult to lyse organisms require stringent lysis steps and low biomass samples require additional rounds of PCR. A host of bioinformatics tools are available for removing residual sequencing errors (18, 45–47). Other tools are available for removing chimeras (14, 35) where there is a trade off between the sensitivity of detecting chimeras and the specificity of correctly calling a sequence a chimera. In recent years, parameters for these algorithms have been changed to increase their sensitivity with little evaluation of the effects on the specificity of the algorithms (45, 47). Others recommend removing any read that has an abundance below a specified threshold as a tool to remove PCR and sequencing artifacts (e.g. removing all sequences that only appear once) (20, 45–47). This method must be approached with caution as such approaches are likely to introduce a different bias of the community representation and ignore the fact, as we showed, that artifacts may be quite abundant and reproducible. Ultimately, researchers must test their hypotheses with multiple methods to validate the claims they reach with any one method (48). All methods have biases and limitations and we must use complementary methods to obtain robust results.

## Materials & Methods

### Mock community

The ZymoBIOMICS™ Microbial Community DNA Standard (Zymo, CA, USA) was used for mock communities and the bacterial component was made up of *Pseudomonas aeruginosa, Escherichia coli, Salmonella enterica, Lactobacillus fermentum, Enterococcus faecalis, Staphylococcus aureus, Listeria monocytogenes*, and *Bacillus subtilis* at equal genomic DNA abundance (https://web.archive.org/web/20171217151108/http://www.zymoresearch.com:80/microbiomics/microbial-standards/zymobiomics-microbial-community-standards). The actual relative abundance for each bacterium was obtained from Zymo’s certificate of analysis for the lot (Lot: ZRC187325), which they determined using shotgun metagenomic sequencing (https://github.com/SchlossLab/Sze_PCRSeqEffects_mSphere_2019/data/references/ZRC187325.pdf).

### Human samples

Fecal samples were obtained from 4 individuals who were part of an earlier study (49). These samples were collected using a protocol approved by the University of Michigan Institutional Review Board. For this study, the samples were de-identified. DNA was extracted from the fecal samples using the MOBIO™ PowerMag Microbiome RNA/DNA extraction kit (now Qiagen, MD, USA).

### PCR protocol

Five high fidelity DNA polymerases were tested including AccuPrime™ (ThermoFisher, MA, USA), KAPA HIFI (Roche, IN, USA), Phusion (New England Biolabs, MA, USA), Platinum (ThermoFisher, MA, USA), and Q5 (New England Biolabs, MA, USA). Manufacturer recommendations were followed except for the annealing and extension times, which were selected based on previously published protocols (18, 38). Primers targeting the V4 region of the 16S rRNA gene were used with modifications to generate MiSeq amplicon libraries (18) (https://github.com/SchlossLab/MiSeq_WetLab_SOP/). The 16S rRNA gene targeting regions of the primers annealed to *E. coli* positions 515 to 533 (GTGCCAGCMGCCGCGGTAA) and 787 to 806 (GGACTACHVGGGTWTCTAAT). The number of rounds of PCR used for each sample and polymerase started at 15 and increased by 5 rounds up to 35 cycles. Insufficient PCR product was generated using 15 rounds and has not been included in our analysis.

### Library generation and sequencing

Each PCR condition (i.e. combination of polymerase and number of rounds of PCR) were replicated four times for the mock community and one time for each fecal sample. Libraries were generated as previously described (18) (https://github.com/SchlossLab/MiSeq_WetLab_SOP/). The libraries were sequenced using the Illumina MiSeq sequencing platform to generate paired 250-nt reads.

### Sequence processing

The mothur software program (v 1.41) was used for all sequence processing steps (50). The protocol has been previously published (18) (https://www.mothur.org/wiki/MiSeq_SOP). Briefly, paired reads were assembled using mothur’s make.contigs command to correct errors introduced by sequencing (18). Any assembled contigs that contained an ambiguous base call, mapped to the incorrect region of the 16S rRNA gene, or appeared to be a contaminant were removed from subsequent analyses. Sequences were further denoised using mothur’s pre.cluster command to merge the counts of sequences that were within 2 nt of a more abundant sequence. The VSEARCH implementation of UCHIME was used to screen for chimeras (35, 51). At various stages in the sequence processing pipeline for the mock community data, the mothur seq.error command was used to quantify the sequencing error rate as well as the true chimera rate. This command uses the true sequences from the mock community to generate all possible chimeras and removes any contigs that were at least three bases more similar to a chimera than to a reference sequence. The command then counts the number of substitutions, insertions, and deletions in the contig relative to the reference sequence and reports the error rate without the inclusion of chimeric sequences (19). UCHIME’s sensitivity was calculated as the percentage of true chimeras that were detected as chimeras when using UCHIME. Its specificity was calculated as the percentage of non-chimeric sequences that were detected as being non-chimeric by UCHIME. The reference sequences and *rrn* operon copy number for each bacterium were obtained from the ZymoBIOMICS™ Microbial Community DNA Standard protocol (https://web.archive.org/web/20181221151905/https://www.zymoresearch.com/media/amasty/amfile/attach/_D6305_D6306_ZymoBIOMICS_Microbial_Community_DNA_Standard_v1.1.3.pdf). Sequences were assigned to operational taxonomic units (OTUs) at a threshold of 3% dissimilarity using the OptiClust algorithm (52). To adjust for unequal sequencing when measuring alpha and beta diversity, all samples were rarefied for downstream analysis. The Good’s coverage for the samples was routinely greater than 95%.

### Statistical analysis

All analysis was done with the R (v 3.5.1) software package (53). Data transformation and graphing were completed using the tidyverse package (v 1.2.1). The distance matrix data was analyzed using the adonis function within the vegan package (v 2.5.4).

### Reproducible methods

The data analysis code for this study can be found at https://github.com/SchlossLab/Sze_PCRSeqEffects_mSphere_2019. The raw sequences are available at the SRA (Accession SRP132931).

## Acknowledgements

We appreciate the willingness of the donors to provide stool samples. We also thank Judy Opp and April Cockburn for their assistance in sequencing the samples as part of the Microbiome Core Facility at the University of Michigan. Additional thanks to members of the Schloss lab and Dr. Marcy Balunas for reading earlier drafts of the manuscript and providing helpful critiques. Support for MAS came from the Canadian Institute of Health Research and NIH grant UL1TR002240 and support for PDS came from NIH grants P30DK034933, R01CA215574, and U19AI09087.

---

**Figure S1: With the exception of the sequence data generated using the KAPA polymerase, the ratio of the two *Salmonella enterica* V4 sequences from the mock community was lower than the expected ratio of 6:1.** The dominant and rare *S. enterica* V4 sequences differed by a single base. The horizontal gray line indicates the expected 6:1 ratio. Each line represents the mean of four replicates.

