## Supplementary figures and images for "The impact of DNA polymerase and number of rounds of amplification in PCR on 16S rRNA gene sequence data"

### Figure S1

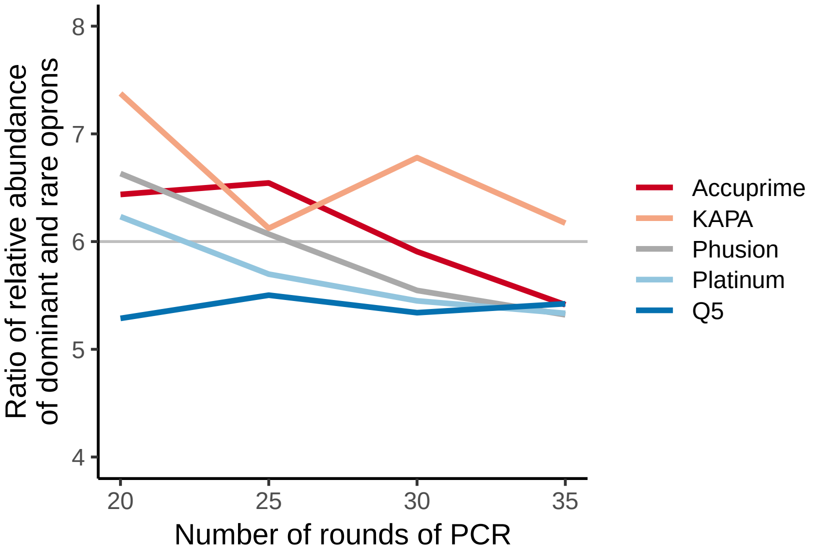
